# MINSTED nanoscopy enters the Ångström localization range

**DOI:** 10.1101/2022.03.18.484906

**Authors:** Michael Weber, Henrik von der Emde, Marcel Leutenegger, Philip Gunkel, Volker C. Cordes, Sivakumar Sambandan, Taukeer A. Khan, Jan Keller-Findeisen, Stefan W. Hell

**Affiliations:** Department of NanoBiophotonics, Max Planck Institute for Multidisciplinary Sciences, Am Fassberg 11, Göttingen 37077, Germany; Department of Cellular Logistics, Max Planck Institute for Multidisciplinary Sciences, Am Fassberg 11, Göttingen 37077, Germany; Synaptic Metal Ion Dynamics and Signaling, Max Planck Institute for Multidisciplinary Sciences, Am Fassberg 11, Göttingen 37077, Germany; Laboratory of Neurobiology, Max Planck Institute for Multidisciplinary Sciences, Am Fassberg 11, Göttingen 37077, Germany; Department of Optical Nanoscopy, Max Planck Institute for Medical Research, Jahnstraße 29, Heidelberg 69028, Germany

## Abstract

We report all-optical, room-temperature localization of fluorophores with precision in the Ångström range. These precisions are attained in a STED microscope, by encircling the fluorophore with the low-intensity edge of the STED donut beam, while constantly increasing the absolute donut power. Individual fluorophores bound to a DNA strand are localized with *σ* = 4.7 Å, corresponding to a fraction of the fluorophore size, with only 2,000 detected photons. MINSTED fluorescence nanoscopy with single-digit nanometer resolution is exemplified by imaging nuclear pore complexes and the distribution of nuclear lamin in mammalian cells labeled by transient DNA hybridization. Since our experiments yield a localization precision *σ* = 2.3 Å, estimated for 10,000 detected photons, we anticipate that MINSTED will open up entirely new areas of application in the study of macromolecular complexes in cells.

---

Since the 1970s, fluorescence microscopy has been indispensable for studying the distribution of biomolecules in cells. At the turn of this century, stimulated emission depletion (STED) microscopy^1^ broke the diffraction barrier that imposed an apparently unsurmountable physical limit on optical resolution, opening up the imaging of cells at the tens of nanometers scale. This transformation has become possible by relying on a new principle for discerning fluorophores at subdiffraction distances, namely the on/off switching of the ability of fluorophores to fluoresce. The recently introduced MINFLUX^2^ and MINSTED^3^ nanoscopy added another factor of ten, thus finally reaching a resolution at the scale of the fluorescence labels.

MINFLUX and MINSTED uniquely combine the specific strongpoints of STED^4^ and the method called PALM^5^/STORM^6^. Like the latter, they switch the fluorescence ability individually per fluorophore, ensuring the finest discrimination possible of neighboring fluorophores. However, unlike in PALM/STORM where the stochastic, initially unknown position of the fluorophore is derived from the diffraction spot of fluorescence detections emerging on a camera, in MINFLUX and MINSTED the individual fluorophores are localized with a movable reference point in the sample, that is usually defined by the intensity minimum of a donut-shaped beam. By moving this donut minimum closer to the position of the fluorophore during the localization process, MINFLUX and MINSTED increase the information gain per detected photon so that precisions of σ ≈ 1-2 nm are routinely attained with only 200-1,000 photons on single fluorophores. Clearly, once 3*σ* < 1 nm and the molecular construct linking the fluorophores to the target biomolecules is controlled, structural biology type of studies inside cells should become viable using optical microscopes.

A major factor limiting the attainable precision is the background, i.e. photon detections not stemming from the target fluorophore. Initial experiments have shown that MINSTED has an advantage over MINFLUX in this regard, because its donut-shaped STED beam is designed to suppress fluorescence. This is contrary to MINFLUX where the donut elicits fluorescence in an area that is about three times larger than in a standard confocal microscope. Besides, building on a STED microscope that inherently offers resolution tuning by changing the donut power, a MINSTED setup can readily accommodate a resolution ranging from the diffraction limit to the molecular scale. Nonetheless, our initial MINSTED study revealed that subtle heating by the STED beam, probably of the sample and the lens immersion oil, limits the precision to *σ* > 1 nm. This is because the popular STED beam of wavelength *λ*_STED_ = 775 nm entails a several orders of magnitude higher average power than what is typically used in confocal and MINFLUX microscopy. By and large, if cryogenic temperatures^7^ are not acceptable, finding a solution that further reduces *σ* is exceedingly challenging.

Here, we report MINSTED attaining all-optical fluorophore localization with precisions in the Ångström range. Corresponding to a fraction of the fluorophore size, these precisions are attained at room temperature using a STED microscope. Individual fluorophores on a DNA strand are localized with *σ* = 4.7 Å, measured by deriving localizations from arbitrarily chosen blocks of 2,000 photons of single emission traces. For the total of 10,000 photons actually detected in the traces, a precision *σ* = 2.3 Å is derived. MINSTED fluorescence nanoscopy with nanometer resolution is exemplified by imaging nuclear pore complexes in mammalian cells labeled by DNA hybridization as in the method called DNA PAINT^8^. Similar resolution is obtained in MINSTED images of the distribution of synaptic proteins in rat hippocampal neurons. These advancements have become possible because, unlike standard STED, MINSTED nanoscopy operates with just a single on-state fluorophore at a time, whilst all other fluorophores in the focal region are off. Moreover, having just a single active fluorophore enables a more effective implementation of STED to the benefit of the MINSTED concept.

The physics behind our study can be outlined as follows (Fig. 1). In virtually all STED microscopes, including in our initial MINSTED implementation, *λ*_STED_ is tuned to the very red edge of the fluorescence spectrum. For red-orange emitting fluorophores, the popular near-infrared *λ*_STED_ = 775 nm is typically chosen. The reason is that at room temperature, the excitation spectrum of most fluorophores extends deeply into the emission peak. STED donuts with shorter *λ*_STED_ therefore tend to ‘directly’ excite many bystander fluorophores in the anti-Stokes mode, overall producing significant fluorescence in the donut region. This fluorescence consequently compromises the on/off contrast needed for fluorophore separation^9^ (Fig. 1b,c). Unfortunately, as the fluorophore cross-section *ς* for stimulated emission scales with the emission spectrum, shifting *λ*_STED_ far out to the red edge decreases *ς* and thus the STED efficiency per unit STED beam power. Compensating the decrease in *ς* with increasing power clearly has (thermal) limits that become apparent when localizing on the finest scale.

**Fig. 1.**
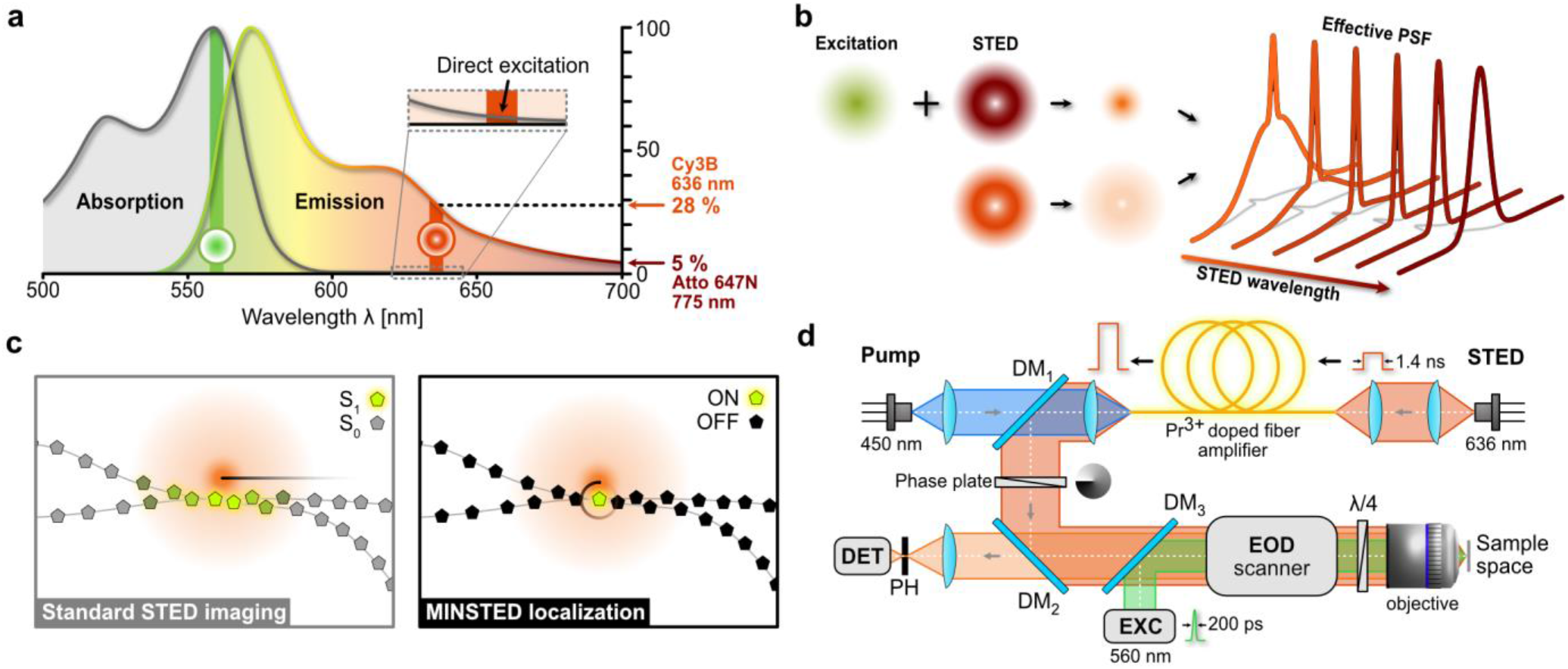
Blue-shifted MINSTED. **a** Qualitative fluorescence and absorption spectra of the fluorophore Cy3B, including our selection of wavelength for excitation (560 nm, green) and de-excitation by stimulated emission (636 nm, red). Reaching well into the emission peak, the cross-section for stimulated emission amounts to 28% of its global maximum, at the expense of slight ‘direct’ excitation of ground state Cy3B fluorophores by the donut-shaped STED beam (inset). **b** Blue-shifting the wavelength of the donut (lower donut has shorter wavelength) for a given power sharpens the central peak of the Effective PSF of the STED microscope but gives rise to a pedestal. **c** The pedestal leads to weak fluorescence from bystander fluorophores, thus compromising the contrast in standard STED imaging (left). Since only one fluorophore is active in MINSTED, the pedestal is ineffectual (right), meaning that the benefits of the blue-shifted STED wavelength can be exploited. **d** Schematic of the MINSTED setup: originating from a 636 nm emitting laser diode, the STED 1.4 ns pulses are amplified by a Pr^3+^ doped fiber pumped with 450 nm laser diode, deflected by a dichroic mirror (DM1), converted into a donut by a phase-plate and aligned with a laser emitting 200 ps pulses for excitation at 560 nm. The co-aligned beams are steered in the focal plane of the objective lens by an electro-optical deflector (EOD), while the quarter-wave plate (*λ*/4) ensures circular polarization. Fluorescence collected from the sample is de-scanned, spatially filtered by a pinhole (PH) and detected.

Fortunately, when the on/off contrast is provided by a process other than STED, as it is the case when switching single fluorophores between active and inactive states, *λ*_STED_ can be tuned closer to the emission maximum so that *ς* becomes larger. The background due to ‘direct’ excitation by the STED donut beam is not of concern in this case, because all fluorophores, apart from the one to be localized, are inactive (Fig. 1c). The larger *ς* enables a lower STED beam power so that prohibitive heating can be avoided and pulsed diode lasers can be used (Fig. 1d).

Blue-shifting *λ*_STED_ also changes the effective point-spread-function (E-PSF) of the optical setup (Fig. 1b). Being the product of the normalized probability for excitation at *λ*_EXC_ (and *λ*_STED_) and that for de-excitation at *λ*_STED_, the E-PSF represents the probability of a fluorophore to emit at a certain coordinate in the focal region. Generally, the full-width-half-maximum (FWHM) of the E-PSF becomes narrower with increasing *ς* and donut intensity. On the other hand, the blue-shifted *λ*_STED_ causes a pedestal due to ‘direct’ excitation (Fig. 1b). While this pedestal compromises bulk STED imaging (Fig. 2a,b), it is not of concern when addressing solitary emitters.

**Fig. 2.**
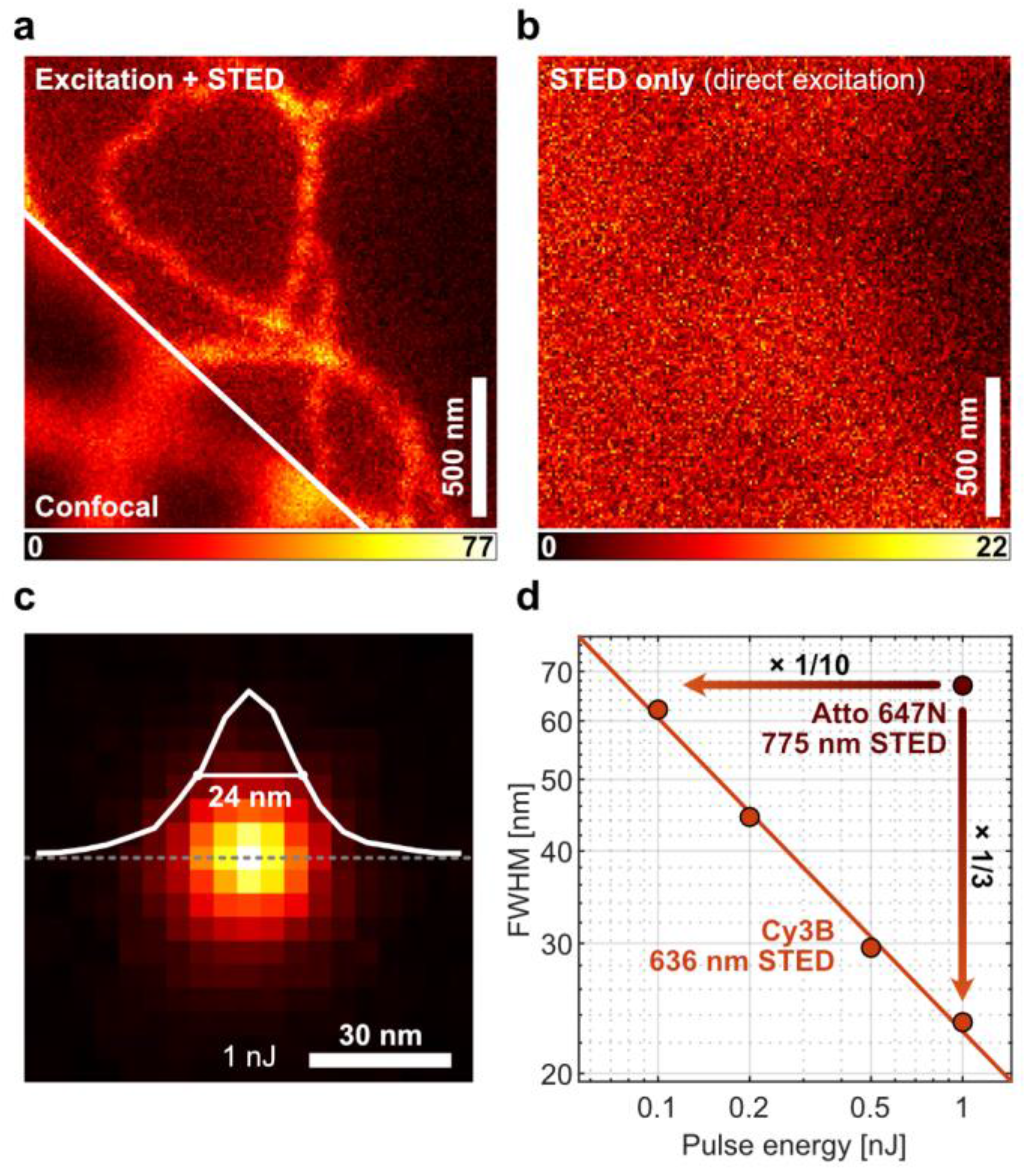
Contrast and resolution of blue-shifted STED. **a** Confocal and STED comparison images of cellular vimentin immunolabeled with Cy3B using 636 nm wavelength for STED. Note the haze around the vimentin fiber images due to the E-PSF pedestal. **b** Corresponding fluorescence image produced by ‘direct’ excitation with the STED beam. In both **a** and **b**, a STED pulse energy of 0.5 nJ was applied. **c** E-PSF with central profile and 24 nm FWHM at STED pulse energy E = 1 nJ, measured with immobilized single Cy3B molecules. **d** FWHM of E-PSF as a function of E; FWHM measured with standard 775 nm STED beam on Atto 647N molecules is displayed for comparison.

For the fluorophore Cy3B with emission peaking at ≈ 570 nm we implemented *λ*_STED_ = 636 nm so that *ς* reached 28 % of its global maximum. This should be contrasted with the 5% obtained when applying the popular *λ*_STED_ = 775 nm to red-emitting fluorophores like Atto647N. The co-aligned *λ*_EXC_ = 560 nm excitation and STED beams, having electronically synchronized pulses of 200 ps and 1.4 ns duration, respectively, were focused into the sample by an oil-immersion objective of 1.4 numerical aperture. Thus, an E-PSF of 24 nm lateral FWHM was gained at a STED pulse energy of 1 nJ (Fig. 2c,d). In comparison, pulses of ≈ 10 nJ would be needed to attain the same FWHM with Atto647N at *λ*_STED_ = 775 nm. At the used 10 MHz repetition rate, this blue-shift entailed a reduction of the average STED beam power from 100 mW to 10 mW, thus substantially lowering the thermal load. Note that tuning the FWHM continually from confocal down to a minimal FWHM is an integral part of each MINSTED fluorophore localization.

For separation by on/off switching, we first opted for the mechanism implemented in the method called DNA PAINT: the transient binding of fluorophores to the biomolecular targets of interest via DNA hybridization. The fluorophore is ‘on’ when, bound to a target, it emits from the same coordinate. Conversely, the fluorophore is ‘off’ when it diffuses in the surrounding medium, generating just a weak background (Fig. 3). This on/off modulation by binding and diffusion allowed us to avoid photoactivatable and -switchable fluorophores and employ regular fluorophores instead, specifically Cy3B. In general, the combination of STED with labeling by DNA hybridization is highly synergistic, because when localizing a bound fluorophore, the STED donut suppresses the background from the diffusing fluorophores. The use of STED simply increases the DNA labeling contrast by adding another off-switching mechanism. Likewise, this amplified off-switching facilitates employing higher concentrations of diffusing labels compared to standard high-resolution DNA PAINT applications, so that the imaging can be accelerated^10^.

**Fig. 3.**
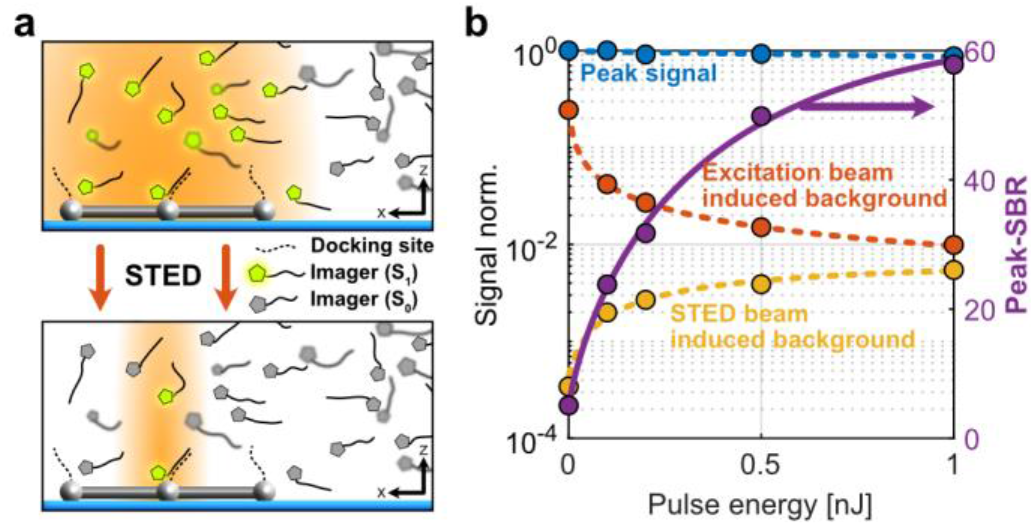
Combining STED with on/off switching and labeling by DNA hybridization. **a** Fluorophores (pentagons in grey and highlighted in green when able to fluoresce) attached to single stranded DNA diffusing in solution, sporadically binding to molecular targets having complementary DNA strands; here the target is a DNA origami represented by grey spheres and sticks. The region in which fluorescence is possible (i.e. E-PSF region) is shown in orange for the confocal (top panel) and the STED case (lower panel). Suppression of the fluorescence of the quickly diffusing fluorophores by STED increases the ratio between the fluorescence signal of bound (on) and diffusing (off) fluorophores. The increased on/off ratio enhances the detection of single bound fluorophores. Conversely, it can be used to increase the concentration of diffusing fluorophores so to increase the imaging speed. **b** Peak fluorescence from single DNA-bound Cy3B fluorophores (blue), fluorescence from diffusing fluorophores in STED single fluorophore recording (red), and STED beam induced fluorescence (orange) as a function of the STED pulse energy E. The signal to background ratio (SBR) increases by a factor > 10 over that of confocal microscopy due to application of E = 1 nJ STED pulses.

To quantify our blue-shifted MINSTED localization and nanoscopy, we carried out measurements with rectangular DNA origami arrays offering 3×3 binding sites for individual fluorophores at 12 nm periodic distance. The concentration of diffusing Cy3B fluorophores was chosen such that only one fluorophore docked to the grid points within the focal region at a time, while the other fluorophores diffused freely in solution (Figs. 3b and 4a-c). First, we verified the gain in signal to background ratio (SBR) with increasing STED pulse energy *E* (Fig. 3b). To this end, we raster-scanned a 2 μm × 2 μm field of view, applying an excitation average power of 1.5 μW. The peak fluorescence rendered by individually bound fluorophores was extracted from the fluorescence maxima in the resulting image, while the background was derived from the mean fluorescence signal per pixel. Recordings with the excitation beam turned off allowed us to quantify the ‘direct’ excitation by the STED beam. We found that the dominant background component was indeed due to the excitation beam inducing fluorescence from diffusing fluorophores. However, this component rapidly dropped with increasing STED pulse energy *E*, since the donut confined the region where fluorescence was allowed. ‘Direct’ excitation by the STED beam increased with *E*, but remained acceptable (Fig. 3b). Altogether, the resulting SBR = 60 at 1 nJ was > 10 times higher than the SBR obtained by standard confocal microscopy (*E* = 0 nJ), and also not sacrificed as in the typical last localization steps of MINFLUX, setting the ground for high-precision localization.

Localization of a fluorophore docking onto a binding site was accomplished by scanning the co-aligned excitation and STED beams circularly around the fluorophore such that, while continuously increasing *E*, it always experienced the steep edge of the E-PSF, i.e. the region where the probabilities of spontaneous and stimulated emission are roughly equal. This condition was kept by adjusting the scanning radius *R* to half of the FWHM of the E-PSF. Thus, the detection probability became indicative of the fluorophore’s position with respect to the donut zero and the fluorophore always experienced the same low intensity of the STED beam. While we started out with a confocal E-PSF, *E* was increased for every detected photon *i*, while maintaining the *R*_*i*_ = FWHM_*i*_/2 condition. At the same time, the circle center was shifted towards the direction of each detection by 0.15 *R*_*i*_. After reaching *E* = 1 nJ and the minimal radius *R*_min_= 24 nm / 2 = 12 nm, which was typically the case for *i* = *N*_*c*_ ≈ 80, the E-PSF and *R*_*i*_ were left constant whereas the center position was still updated until the localization ended. The resulting measurement is a series of circle center positions (*x*_*j,i*_, *y*_*j,i*_) representing fluorophore coordinate updates until *i* = *L*. The index *j* refers to different localizations, i.e. fluorophores. We recorded up to *j* = 991 localizations, spread over ≈ 144 binding sites. This set of center positions was used to quantify the lateral localization precision of the nanoscope.

Of all localizations, 90 % featured a standard deviation 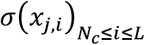 of the center positions below *σ* _c_ = 4.2 nm in both x- and y-direction. In other words, once the minimal FWHM at about *i* = *N*_*c*_ was reached, the localizations converged within an area defined by *σ* _c_. The remaining 10% were compromised by the binding of a second fluorophore or other sources of background. The vast majority of localizations could be assigned to specific binding sites of the grid patterns just by grouping the localizations in clusters maximally covering 10 nm in diameter. Clusters with less than five localizations were discarded. Provided the binding sites are firm, the multiple localizations per cluster yield the localization precision as a function of the number of detections *N*. For 1 < *N* ≤ *N*_c_, the precision *σ* _cluster_(*N*) from the cluster analysis was established as the standard deviation 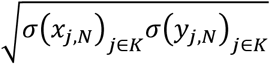 of the center positions (*x*_*j,N*_, *y*_*j,N*_) in each cluster of index *K*. Once the minimum FWHM had been reached, i.e. for *N* > *N*_c_, the fluorophore position was equated with the average of the measured center positions 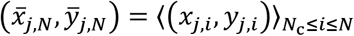. Again, the standard deviation of these values was taken as the resulting precision *σ* _cluster_(*N*). Plotting *σ* _*c*luster_ as a function of *N* shows that in the range of continuously decreasing FWHM, i.e. for 1 < *N* < *N*_c_, the median cluster spread rapidly scaled down to about 4 nm at *N* = *N*_c_ (Fig. 4a). For *N* > *N*_c_, *σ* _cluster_ improved more slowly because it scaled just with 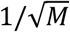, with *M* = *N* − *N*_c_ + 1 being the number of detections after the minimal FWHM had been reached. For *N* > 1,000, *σ* _cluster_ levels off at slightly below 1 nm, probably due to residual drift of the binding sites. Note that the measure *σ* _cluster_ includes, besides the pure MINSTED localization precision, all thermal and mechanical disturbances of both the microscope and the sample over the whole course of the experiment lasting for 40 min at room temperature.

**Fig. 4.**
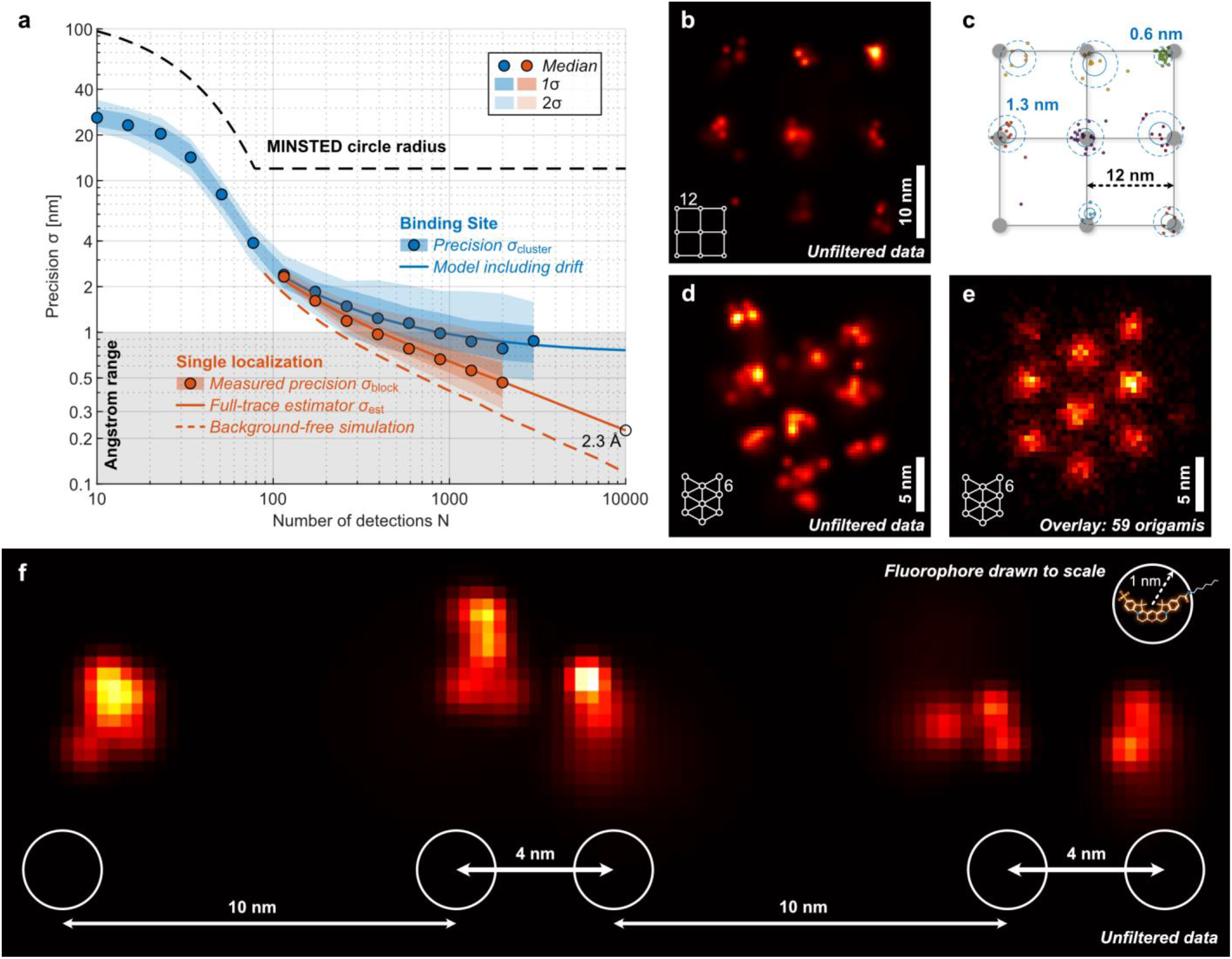
Localization precision and resolution in MINSTED nanoscopy. **a** Localization precision (points: median; shaded areas: ±1 and ±2 standard deviations) measured from many consecutive binding events on clustered binding sites (blue) on DNA origami grids of 12 nm periodicity and for each binding event individually (red). Blue points and shades are only displayed if computed from at least ten clusters. The red solid line shows the estimated localization precision for the individual events; resulting from that, instabilities are considered to reconstruct the cluster data (blue solid line). Simulated localizations without background are shown as red dashed line. **b** MINSTED image of rectangular binding site pattern of 12 nm periodicity and the pertinent localization distribution in **c**. Each localization is represented by its estimated position. Blue circles correspond to one (solid) and two (dashed) standard deviations of the estimated binding site position. The cluster identity is color coded. **d** MINSTED image of 3×3 hexagonal DNA origami with internal distances of 6 nm. **e** Overlay of 59 MINSTED images of the origami pattern from **d**, completely resolving the periodically arranged bindings of 6 nm mutual distance. **f** Binding sites of 4 nm distance are fully resolved by MINSTED; sketch of the underlying origami design is shown below. The circles of 2 nm diameter represent the extent of the Cy3B molecules whose structure is drawn to scale (upper right corner), in order to highlight the relationship between the localization precision and the fluorohore size.

Since a single localization took only ≈ 200 ms, estimating the localization precision on the basis of a single localization allowed us to contain potential influences of movements. For each localization with *L* − *N*_c_ + 1 detections at *R*_min_, the localization precision for *i* ≥ *N*_c_ was obtained by calculating the standard deviation *σ* _block_(*M*) of a moving mean of overlapping blocks of *M* center positions (*x*_*i*_, *y*_*i*_). To ensure at least five independent data blocks, only blocks of size *M* < (*L* − *N*_*c*_ + 1)/5 were considered. To quantify single localization trace precisions up to *M* = *L* − *N*_*c*_ + 1, the values of *σ* _blo*c*k_(*M*) for *M* = 1, 2, …, ⌊(*L* − *N*_c_ + 1)/5⌋ were fitted to a power-law model *σ* _est_(*M*) = *a*/(*b* + *M*)^*c*^ with parameters *a, b, c*. The parameter *b* > 0 accounts for correlations among short time spans caused by the fractional updates of the center positions, while *c* ∈ (0.4, 0.5] allows for the non-ideal use of photon information. To keep the two estimates *σ* _blo*c*k_(*M*) and *σ* _est_(*M*) comparable within the whole range of photon numbers shown, only traces with *L* − *N*_*c*_ + 1 ≥ 10,000, amounting to 39 traces in total, are displayed. At *N* = *M* + *N*_*c*_ − 1 = 2,000 detected photons, a median localization precision of 4.7 Å was obtained from both estimators, which clearly indicates the ability of MINSTED to localize at a fraction of the fluorophore’s size of about 2 nm. The precision of 1 nm is attained with a new record of only 400 detections. With 10,000 detections in place and under the same realistic background and stability conditions, the estimated precision *σ* _est_ improved to 2.3 Å (Fig. 4a).

Comparing the single localization estimate *σ* _est_ and the cluster analysis precision *σ* _cluster_ enabled us to assess the effective position uncertainty *s* of the binding sites over the 40 minutes of measurement as a proxy for the stability implicated in the process. By modeling 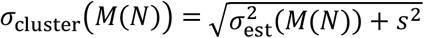, we obtained a value of *s* = 0.73 nm, which proves the long-term stability of our system. The combination of precision and stability enabled our blue-shifted MINSTED system to clearly resolve origami binding sites as close as 4 nm, which is about twice the molecular size of Cy3B (Fig. 4b-f). Registering all localizations with *L* > *N*_*c*_ resolved the entire origami pattern.

To explore the performance of blue-shifted MINSTED nanoscopy in biological samples, we prepared mammalian (HeLa) cells expressing the nuclear pore protein NUP96 as polypeptides carboxy-terminally tagged with sfGFP serving as binding sites for anti-GFP-nanobodies. Due to the NPCs’ ideally eightfold symmetry in the focal plane, NPC scaffold components like NUP96 are frequently used to assess nanoscopy methods^11^. The nanobodies targeting the GFP tags of these NUP96 polypeptides carried a DNA-docking strand to which a complementary DNA strand with a Cy3B fluorophore was able to dock by hybridization. About 30% of the localization attempts did not converge and were discarded. The remaining 70 % rendered images with a median *σ* < 1 nm (Fig. 5a-d). Since the fluorescence images yield only the fluorophore distribution, they must be seen as a proxy of the actual NUP96 distribution in the cell with the 3*σ* = 5-7 nm uncertainty given by the size of the GFP-nanobody construct. This should be contrasted with our 3*σ* < 3 nm for the fluorophore localization in the cells, proving that in current fluorescence nanoscopy the main limits for extracting positional information of the biomolecules is the size of the tags. Nonetheless, creating an overlay image by averaging over only 77 pores consisting of 2,006 localizations nicely reproduced the NPC’s eightfold symmetry as visualized via NUP96 (Fig. 5f-h). The elongation of each NUP96 localization cluster along the periphery in the image can be interpreted as being indicative of the slightly staggered arrangement of the NUP96 polypeptides within each of the NPC’s octagonal subunits^12^. After taking the linker length into account, the average diameter of 111 nm as extracted from our data (Fig. 5g) is in excellent agreement with results based on electron microscopy^13^.

**Fig. 5.**
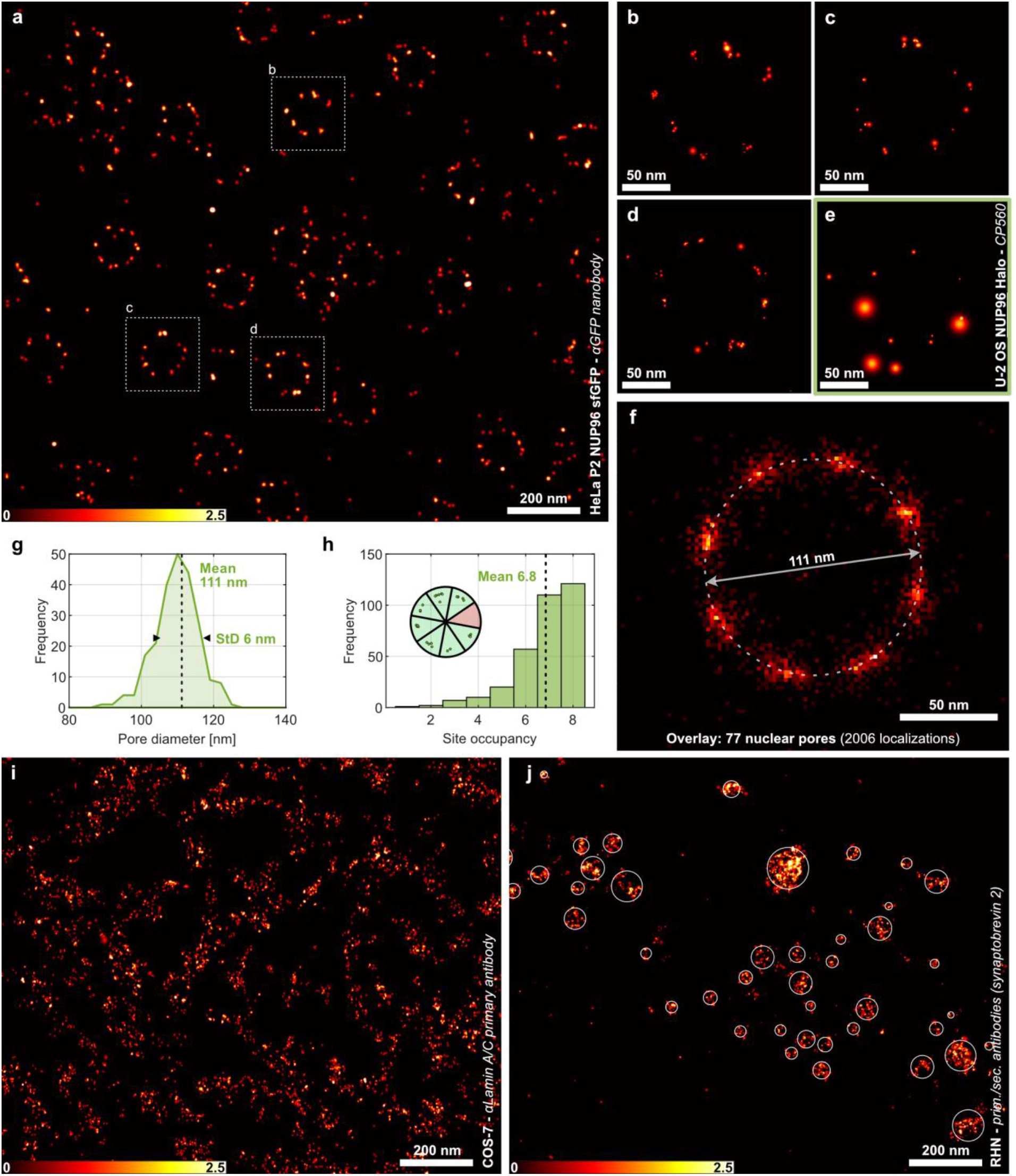
Blue-shifted MINSTED imaging in cells. **a** MINSTED image of the nuclear surface of a HeLa P2 cell expressing nuclear pore protein NUP96 endogenously tagged with sfGFP and labelled with nanobody against GFP offering a DNA binding site for hybridization with a complementary DNA strand labeled with the fluorophore Cy3B. **b-d** Excerpts of individual nuclear pore images from panel **a**, as indicated in the boxed region highlighting the median localization precision of 0.9 nm. As NUP96 occurs in 4 copies per one-eighth of the eightfold rotationally symmetric NPC, each ‘corner’ is expected to harbor up to four binding sites, which agrees well with the several individual dots in the images. **e** NUP96 image from a U-2 OS cell recorded by Halo-tag labeling with the photoactivatable fluorophore Halo-ONB-CP560. While this labeling does not attain the same level of completeness as DNA hybridization, the image highlights the viability of blue-shifted MINSTED with photoactivatable fluorophores offering just a single on-off cycle. **f** Overlay image of 77 HeLa nuclear pore MINSTED images renders the expected eightfold symmetry of the nuclear pores, with each corner displaying an elongation along the circumference, indicative of the slightly staggered arrangement of the NUP96 polypeptides adjacent to each other in each of the NPC’s octagonal subunits. **g** The average diameter as formed by NUP96 is 111 nm. **h** The number of occupied subunits within the NUP96 eight-fold-symmetry displays a mean of 6.8. **i** Imaging of lamin A/C exemplifies the application of MINSTED at high labelling densities. **j** MINSTED image of synaptic vesicles in neurons represented by primary and secondary antibody-tagged synaptobrevin 2. The shown clusters of single fluorophore events yield images of tagged synaptic vesicles that appear as entities of (38 ± 23) nm in diameter (4σ). Except **f**, all images were rendered by displaying the individual localizations by Gaussian functions with standard deviation corresponding to the localization precision.

While labeling by DNA hybridization has many advantages over the use of photoactivation for on/off switching, blue-shifted MINSTED nanoscopy also works with the latter. This is demonstrated by labeling U-2 OS NUP96-Halo cells^11^ with the photoactivatable fluorophore Halo-ONB-CP560 having an absorption and emission maximum at 560 nm and 610 nm, respectively. Since each photoactivatable fluorophore provides just a single localization, the resulting NUP96 image displays a lower localization density (Fig. 5e). Here, a median *σ* of 2 nm was observed, while the median value of *L* − *N*_*c*_ + 1 was reduced to 145. Yet, this example shows that once photoactivatable fluorophores are optimized, precisions as with Cy3B can be attained throughout.

Having passed this test, MINSTED nanoscopy was next applied to structures not exhibiting a similarly symmetric arrangement of its components like the NPCs. As one example, we visualized the nuclear lamina, the filamentous network that occurs positioned between the NPCs at the nuclear side of the nuclear envelope. Specifically, we labeled COS-7 cells with anti-lamin A/C antibodies having two DNA docking strands at their glycosylation sites close to the antibody binding pocket. The docking strands served for Cy3B labeling through DNA hybridization. The antibodies target the Ig-fold domain of lamin A/C. Recording the fluorophores and their positions for 68 minutes yielded a nanoscale proxy of the lamin distribution (Fig. 5i). The high binding site density infrequently caused undue binding of more than one fluorophore in the focal region, which we excluded. Although this sample posed more challenges for separation and localization, the median *σ* was < 1 nm. Finally, we applied MINSTED to the distribution of synaptic vesicles in a cultured rat hippocampal neuron, highlighted through tagging the protein synaptobrevin 2 with a primary/secondary antibody sandwich. The resulting MINSTED image displays fluorophore clusters of (38 ± 23) nm in diameter, in line with the expectation to represent vesicles^14^ (Fig. 5j).

Summarizing our findings, blue-shifted MINSTED is able to localize individual fluorophores with *σ* = 1 nm precision using only 400 detected photons, a record low number. Note that for this precision, an idealized background-free centroid-based localization would require ≈ 11,000 photons, underscoring the benefit of targeting the fluorophore with an intensity (donut) zero. Cutting down the number of required detections also reduces the influence of movements and drift. Fluorophores as close as 4 nm can be fully separated. Apart from discarding failed localizations, no further data post processing or drift correction was required. This success is because i) MINSTED localizes with a STED donut minimum providing a reference coordinate in the sample, which is ii) continuously moved closer to the fluorophore so that the de-excitation rate is kept constant, iii) STED and confocal detection suppress fluorescence background and iv) low power beams can be used to avoid subtle disturbances by heating.

As the 3*σ* value of 3 nm is even smaller than the size of the molecular construct linking the fluorophore to the biomolecule of interest, our results underscore that the limits of fluorescence nanoscopy applications in biology are no longer set by physical or technical factors, but by the molecular linkers. Evidently, a fluorescence image renders just the fluorophores, not the tagged biomolecules. Therefore, further advancements in molecule-scale biological imaging critically call for solutions to relate the position of the fluorophore to that of the target biomolecule. Finding such solutions has become more attractive than ever because MINSTED now enables localizations down to the Ångström domain. In fact, our analysis showed that if 10,000 emissions can be detected from the fluorophore under the same practical background and stability conditions, the precision is estimated to *σ* = 2.3 Å, a value that is about 8 times smaller than the extent of the fluorophore itself. Clearly, once the problem of assigning the fluorophore’s position to that of the labelled target has been solved, such precisions should open up new pathways for studying biomolecular assemblies in cells with optical microscopes under physiological conditions.

Finally, we note that the attained precision records can still be improved without amending the MINSTED concept. As the detection rate is largely proportional to the pulse repetition rate of our diode lasers, increasing the rate from the present 10 MHz to 100 MHz should speed up the localization almost accordingly. As laser technology advances rapidly, this scenario may soon enable MINSTED localization with 1 nm precision in less than about 20 milliseconds, accommodating even faster movements and drifts. Likewise, fluorophore localizations with 1 Å precision should become routine.

## Author contributions

H. v. d. E. and M. W. built the setup and M. L. wrote the software, including the real-time control of the setup. M. W. and H. v. d. E. built the fiber-amplified STED laser system after planning and construction by M.W.. M. W. and H. v. d. E. prepared the samples and H. v. d. E. performed the measurements. T. K. synthesized the photoactivatable dye. H. v. d. E. analyzed the data with feedback from S. W. H., M. W. and M. L., while J. K. F. performed the particle averaging. S. S. prepared the rat hippocampal neuron samples. P. G. generated and characterized the HeLa P2 NUP96-sfGFP cell line supervised by V. C., P.G. prepared the respective samples. S. W. H. outlined the project, initiated and supervised its exploration. S. W. H., M. L., H. v. d. E. and M. W. wrote the manuscript. All authors contributed to the manuscript and the supplementary information either through discussions or directly.

## Funding

This work has been funded by the German Federal Ministry of Education and Research (BMBF) (FKZ 13N14122 to S. W. H). H. v. d. E. is part of the Max Planck School of Photonics supported by BMBF, Max Planck Society, and Fraunhofer Society.

## Acknowledgement

We thank Ellen Rothermel and Tanja Koenen for their help with cell culture and labelling of cells, Vladimir N. Belov for advice regarding the chemical synthesis, the European Molecular Biology Laboratory (Heidelberg) for U-2 OS-CRISPR-NUP96-Halo cells, Nickels Jensen and Christian Brueser for generation and testing of gene edited cell lines and Steffen J. Sahl for the fruitful discussions.

## Competing Interest

The Max Planck Society hold patents on selected procedures and embodiments of MINSTED, benefitting H. v. d. E., M. L., M.W., and S. W. H..

